# Adaptation of the gut holobiont to malnutrition during mouse pregnancy depends on the type of nutritional adversity

**DOI:** 10.1101/2021.10.26.465755

**Authors:** Kristin L Connor, Enrrico Bloise, Todd Z DeSantis, Stephen J Lye

## Abstract

Malnutrition can influence maternal physiology and programme offspring development. Yet, in pregnancy, little is known about how dietary challenges that influence maternal phenotype affect gut structure and function. Emerging evidence suggests that interactions between the environment, multidrug resistance (MDR) transporters and microbes may influence maternal adaptation to pregnancy and regulate fetoplacental development. We hypothesised that the gut holobiont (host and microbes) during pregnancy adapts differently to suboptimal maternal diets, evidenced by changes in the gut microenvironment, morphology, and expression of key protective MDR transporters during pregnancy. Mice were fed a control diet (CON) during pregnancy, or undernourished (UN) by 30% of control intake from gestational day (GD)5.5-18.5, or fed 60% high fat diet (HF) for eight weeks before and during pregnancy. At GD18.5, maternal small intestinal (SI) architecture (H&E), proliferation (Ki67), P-glycoprotein (P-gp - encoded by *Abcb1a/b*) and breast cancer resistance protein (BCRP/*Abcg2*) MDR transporter expression and levels of pro-inflammatory biomarkers were assessed. Circulating inflammatory biomarkers and maternal caecal microbiome composition (G3 PhyloChipTM) were measured. MDR transporter expression was also assessed in fetal gut. HF diet increased maternal SI crypt depth and proinflammatory load, and decreased SI expression of *Abcb1a* mRNA, whilst UN increased SI villi proliferation and *Abcb1a*, but decreased *Abcg2*, mRNA expression. There were significant associations between *Abcb1a* and *Abcg2* mRNA levels with relative abundance of specific microbial taxa. Using a systems physiology approach we report that common nutritional adversities provoke adaptations in the pregnancy holobiont in mice, and reveal new mechanisms that could influence reproductive outcomes and fetal development.

## Introduction

Undernutrition and overnutrition before and during pregnancy adversely impact reproductive outcomes. Both forms of malnutrition are associated with maladaptations to pregnancy, including altered endocrine function[1–3], increased maternal susceptibility to infections and low-grade inflammation[4–6], as well as poor development of the placenta and offspring antenatally and postnatally[7, 8]. Rates of maternal undernutrition and underweight are increasing[8], and progressively, more populations are facing overnutrition and epidemics of obesity[9, 10], including those undergoing transitions in nutrition[11].

In the non-pregnant state, malnutrition can have profound effects on gut structure and function. This includes altered intestinal architecture and histomorphological changes in key cell types[12], and functional changes including reduced barrier integrity/defence and an altered gut microbiome[13–15]. These changes are associated with compromised health outcomes such as poor growth and metabolic/immune dysfunction. The intestinal epithelium directly senses and responds to the external environment[16, 17], and these adaptations can determine allostasis and homeostasis locally at the gut and systemically beyond the gut, collectively to protect the host and influence health. Multidrug resistance (MDR) transporters, such as P-glycoprotein (P-gp - encoded by *Abcb1a/b* genes) and breast cancer resistance protein (BCRP/*Abcg2*), belong to the ATP-binding cassette (ABC) family of transporters, and establish an additional key defensive barrier between the gut and the intestinal lumen that contributes to intestinal homeostasis[18]. MDRs are mostly distributed on the apical surface of intestinal epithelial cells[19–21] and support the physical, selective barrier of the intestinal epithelium through their ability to efflux an array of substances across biological membranes, preventing the gut from absorbing potentially harmful substances into the body. Intestinal P-gp and BCRP regulate gut absorption, biodistribution, and metabolism of an array of obstetric-relevant substrates including: 1. nutrients (folate, haem/iron, flavonoids), 2. cytokines and chemokines (chemokine [C-C motif] ligand 2 [CCL2], granulocyte-macrophage colony-stimulating factor [GM-CSF], interleukin[IL]-1 β, IL-2, IL-4, IL-6, interferon[IFN]-γ, tumor necrosis factor [TNF]-α), 3. toxicants (carcinogens phototoxic compounds, estrogenic mycotoxins, select mercuric species, ivermectin) and 4. drugs (anti-retrovirals, antibiotics, synthetic glucocorticoids, sulfonylureas, nonsteroidal anti-inflammatory drugs, proton pump inhibitors)[22, 23]. Malnutrition[24] and metabolic disease[25, 26] in the non-pregnant state alter MDR expression and function. Yet, little is known about how dietary challenges that influence pregnancy phenotype affect gut structure and function, including gut MDR transporters, in pregnancy.

Importantly the host is not, and probably has never been, an “autonomous entity”[27]. More accurately, there is clear cooperation and dependence between the host and the community of microbes living in and on it. This holobiont (“the unit of biological organisation composed of a host and its microbes”[27]) senses and responds to the environment, and influences host physiology (including states of health and disease)[28, 29] and behaviour[30], affecting an individual’s phenotype. Emerging evidence suggests that one important interaction within the holobiont occurs between gut microbes and intestinal epithelial MDR transporters, where variation in this relationship can lead to an oscillation between health and diseased states[18]. In reproduction, the holobiont is especially important, since this real-time sensor of the pregnancy environment could instruct maternal adaptations to pregnancy, and even regulate fetoplacental development and function[31, 32], possibly through interactions with MDRs.

Although we know how some nutritional exposures affect selected features of the gut[33], including in pregnancy[34, 35], little is known about the interactions between gut microbes and MDR transporters, and what, if any, functional consequence there might be for the host if these relationships are altered. Not all pregnancies exposed to malnutrition show compromised maternal physiology or health, which could be explained by the extent to which the gut holobiont can adapt to changes in the pregnancy environment, and thus act as a selective barrier for trophic factors and xenobiotics[36]. If we can identify the situations where the holobiont maladapts to environmental exposures, and understand the mechanisms that underlie these maladaptations, we are better positioned to pinpoint the most at-risk pregnancies or especially detrimental early environments, and target these mechanistic pathways to correct holobiont function and prevent poor reproductive health and outcomes.

To address this gap, we evaluated the effects of malnutrition during pregnancy on the gut holobiont in a mouse model. We hypothesised that the gut holobiont in pregnancy adapts differently to suboptimal maternal diets, evidenced by changes in the gut microenvironment, gut morphology, and the expression of key MDR transporters that protect the mother and developing fetus. We also hypothesised that these dietary challenges would programme the same transport pathways in the developing fetal gut. In taking a systems physiology approach, our study reveals a role for malnutrition in influencing the holobiont during pregnancy, with implications for pregnancy health and fetal development.

## Methods

### Animal model

The Animal Care Committee at Mount Sinai Hospital approved all experiments. Our animal model has been described in detail previously, and was established to model pregnancy phenotypes seen with undernutrition/underweight (reduced maternal weight gain with fetal growth restriction, without embryonic lethality or preterm birth) and overweight/obesity (maternal metabolic dysfunction, without fetal growth restriction)[3, 34, 35]. In brief, male and female C57BL/6 mice were obtained from Jackson Laboratories and housed in a single room under constant temperature (25°C and 12:12 light-dark cycle) with free access to food and water. Females were randomly divided into three nutritional groups: i) mice fed a control diet (10.1% kcal as fat [23.4% saturated fat by weight]; 64.5% kcal as carbohydrate; 25.4% kcal as protein; Dustless Precision Pellets S0173, BioServe, Frenchtown, NJ, USA) *ad libitum* before mating and throughout pregnancy (CON, n=7); or ii) mice fed a control diet *ad libitum* before mating and until gestational day (GD) 5.5, and then undernourished (UN) by 30% of control intakes from GD5.5-18.5 (n=7); iii) or mice fed a high fat diet (60% kcal as fat [37.1% saturated fat by weight]; 20% kcal as carbohydrate; 20% kcal as protein; D12492, Research Diets, New Brunswick, NJ, USA) *ad libitum* from 8 weeks before mating and throughout pregnancy (HF, n=8). All breeding males were fed control diet *ad libitum* for the duration of the study. Females in estrus were housed with a male overnight at approximately 10 weeks of age, and mating was confirmed by the presence of a vaginal sperm plug the following morning, at which time (GD0.5) pregnant females were housed individually in cages with free access to water and their respective diets.

### Biospecimen collection and processing

At the end of pregnancy (GD18.5, term = 19 days), dams were killed by cervical dislocation. Immediately following, dams were decapitated and trunk blood was collected into heparin-coated tubes for plasma isolation. Fetuses and placentae were rapidly dissected from the uterus, and tissues were flash frozen in liquid nitrogen and stored at −80C, or fixed in 10% neutral-buffered formalin. Litter characteristics (size, resorptions) were not different between groups[3]. One male and one female fetus from each litter was used for later analyses. Maternal small intestine (SI) was isolated from the gastrointestinal tract as previously described[34, 35]. Maternal SI tissue from the mid portion of the SI (representing the jejunum, a key site for nutrient absorption and transport of other small molecules) and the entire fetal gut were flash frozen in liquid nitrogen and stored at −80°C, or fixed in 10% neutral-buffered formalin. Small intestinal intraepithelial lymphocytes were isolated from SI tissue for measurement of cytokines and chemokines, and DNA was extracted from caecal contents for 16S rRNA profiling, as previously described[35, 37]. The maternal brain was removed from the skull and the cerebellum was dissected off the colliculus inferior[38], and flash frozen in liquid nitrogen and stored at −80°C.

### Histological analyses

Five-micrometre paraffin-embedded maternal SI sections were stained with haematoxylin and eosin (H&E) according to standard protocols to assess intestinal architecture and histological components, or with specific primary and secondary antibodies as described below. To determine if suboptimal maternal nutrition confers architectural changes in the SI that have been documented in cases of starvation[12, 14] and diet-induced inflammation[15, 39, 40] in the non-pregnant state, villus height and crypt depth were assessed as described by Beuling *et al*.[41]. In brief, eight images were randomly captured at 20X magnification along the length of an H&E stained SI section from each dam (Leica DMIL LED inverted microscope, Wetzlar, Germany; and QCapture Pro software, Surrey, BC, Canada). The length of one villi and its corresponding crypt were measured in each image (Image J version 1.51) first by determining the villi-crypt junction and then by measuring the length of the villi and crypt from that junction. A single investigator, blinded to the experimental groups, captured the images and performed the measurements.

Sections from each SI were also stained for Ki67, to determine the relative index of proliferation[42], and P-gp and BCRP, to visualise expression and localisation of these multidrug resistance transporters. For Ki67 staining, sections were first rehydrated and then quenched using 0.3% hydrogen peroxide (Fisher, Toronto, ON, Canada) in methanol for 30 min at room temperature (RT). Sections underwent antigen retrieval first by boiling in 1X unmasking solution (DAKO, Mississauga, ON, Canada) for 30 min and next by boiling in 10 mM sodium citrate solution for 8 min. Sections were blocked using serum-free protein blocking solution (DAKO) for 1 h at RT and incubated overnight at 4°C with 1:100 dilution of mouse anti-Ki67 antibody (Novocastra, Concord, ON, Canada) in antibody diluent (DAKO). Next, a 1:200 dilution of goat biotinylated anti-mouse antibody (Vector Labs, Burlington, ON, Canada) in antibody diluent (DAKO) was applied for 1 h at RT, followed by a 1:2000 dilution of streptavidin-horseradish peroxidase (Invitrogen, Burlington, Canada) in 1× PBS for 1 h at RT. DAB peroxidase substrate (Vector Labs, Burlingame, CA, USA) was applied for 1 min 15 sec to visualise the antibody signals. Between all steps, sections were washed three times with PBS with 0.1% Tween20. Sections were counterstained with Gill’s #1 heamatoxylin. Monoclonal mouse IgG1 antibody (DAKO) served as the negative control. Four of eight images randomly captured SI images at 40X magnification (Leica DMIL LED inverted microscope and QCapture Pro software) were used for measuring relative index of proliferation. First, two villi and their crypts in each of the four images (eight villi and eight crypts in total per dam) were identified for quantification. In each villi and crypt, immunoreactive (ir)-Ki67 positive cells and unstained (negative) cells were counted, and these were summed within a villi and a crypt to obtain the total number of cells in the villus and in the crypt. The relative index of proliferation for each villus and crypt was calculated as the percentage of ir-Ki67 positive cells to total cells. The mean relative index of proliferation across the four villi and crypts for each dam was calculated. Image analysis was conducted by a single observer blinded to the experimental groups.

Staining for P-gp (1:500, D-11 Santa Cruz, Mississauga, Canada) and BCRP (1:200, Calbiochem, Etobicoke, ON, Canada) was conducted as described above with the following changes: sections were quenched using 0.03% hydrogen peroxide in 1X PBS for 30 min at RT; secondary antibody was goat biotinylated anti-mouse antibody (1:200, Vector Labs); and DAB peroxidase substrate incubation time to visualise antibody signals was 30 sec for both P-gp and BCRP. From stained sections, eight images containing villi and crypts were randomly captured at 20X magnification (Leica DMIL LED inverted microscope and QCapture Pro software). Visual assessment of staining in each image was conducted by a single observer blinded to the experimental groups as described previously [34, 43, 44]. Staining was assessed in one villi and its adjacent crypt in each of the eight images in the following four regions: the apical membrane of the villus, the remainder of the same villus (not including apical membrane), the apical membrane of the crypt, and the remainder of the crypt (not including the apical membrane).

### Cytokine analysis from maternal plasma and SI IEL protein

Maternal plasma and SI IEL proteins were assayed for cytokine levels according to manufacturer’s instructions using the Bio-Plex Pro Mouse Cytokine 23-Plex Assay (Bio-Rad, Mississauga, Canada) and the Luminex system (Bio-Rad; software v6.0). SI IEL samples from 19 dams were assayed and plasma samples from 14-19 dams were assayed. Cytokine concentrations are expressed as pg/mL for plasma and pg/mg protein for SI IEL total protein. Cytokines with values above or below the standard curve range were excluded from analyses leaving 19 and 15 cytokines for maternal plasma and SI IEL analyses, respectively.

### RNA isolation and mRNA expression in maternal and fetal tissues

Total RNA was extracted from maternal SI and cerebellum following manufacturer’s protocol (QIAGEN RNeasy Plus Mini Kit, Toronto, Canada) and the Tissue Lyser II (Qiagen, Toronto, Canada) at 30 Hz for 3 minutes. RNA was extracted from fetal gut following manufacturer’s protocol (QIAGEN RNeasy Micro kit, Toronto, Canada) and the Tissue Lyser II at 20 Hz for 2 minutes, twice. RNA quality and quantity were assessed using the Experion StdSens Analysis Kit and LapChip (Bio-Rad, Mississauga, Canada) according to manufacturer’s protocol and by nanodrop. 1 µg RNA was reverse transcribed using 5X iScript Reverse Transcription Supermix (Bio-Rad, Mississauga, Canada).

Real-time qPCR (Bio-Rad CFX384, Hercules, CA, USA) was used to measure expression of *Abcb1a* and *Abcb1b* (encoding P-gp) and *Abcg2* (encoding BCRP) in maternal SI and fetal gut. Expression levels in maternal cerebella were measured as a control to evaluate whether malnutrition targeted MDR expression in a tissue-dependent manner. Primer sequences for genes of interest and three stably expressed reference genes (maternal SI: *Actb* [encoding β-actin], *Tbp* [encoding TATA-box binding protein], *Ywhaz* [encoding 14-3-3 protein zeta/delta]; cerebellum: *Ppia* [peptidylprolyl isomerase A], *Tbp, Ywhaz*; fetal gut: *Actb, Tbp, Ywhaz*) are in Table A.1. A non-template control (NTC; absence of cDNA template) was prepared as a negative control for the PCR reaction. Each real-time PCR reaction mix was prepared with SYBR Green JumpStart Taq ReadyMix (Sigma, Oakville, Canada) and forward and reverse primer mix (3 µM) for each gene. Standard curves for each gene were run. Standards, samples and controls were run in triplicate under the following cycling conditions: 95°C for 20s; 40 cycles of 95°C for 5s and 60°C for 20s and a melt curve at 65°C→95°C with a 0.5°C increment every 5s. Relative gene expression for each sample was calculated using the Cq value of the gene of interest relative to the geometric mean of the reference genes Cq values[45].

### Caecal DNA extraction, 16S rRNA profiling and analysis

In a subset of pregnancies from this cohort (n=5/group), DNA was extracted from frozen caecal contents by bead beating and full-length 16S rRNA gene sequences were amplified from variable regions 1 through 9 from each sample, and hybridised to the PhyloChip™ Array (version G3) as we previously described[35]. Empirical operational taxonomic units (eOTUs) were determined from the dataset and abundance tables for each eOTU were calculated as previously described[35, 46, 47].

### Statistics

Outcome measures were tested for normality and unequal variances (Levene test). Data that were non-normal were transformed to achieve normality, where possible. Differences between dietary groups for outcome measures were determined by ANOVA with Tukey’s post hoc, or Welch ANOVA with Games–Howell post hoc, or Kruskal–Wallis test with Steel–Dwass for nonparametric data (p<0.05). Data are presented as means ± standard deviation (SD) or median and interquartile range (IQR), with 95% confidence diamonds in figures. Data transformed for analyses are presented as untransformed values. Relationships between gene expression levels and relative index of proliferation in SI within the whole cohort were determined by Spearman’s correlation with data presented as means ± SD with Spearman’s rho (p<0.05), and 95% density ellipses in figures.

### Analysis of relationships between the microbiome and multidrug resistance transporters

Significant differences between MDR gene expression levels and the microbiome were determined using the Adonis test. Spearman rank correlations test was used to determine relationships between each eOTU’s abundance value and *Abcb1a* and *Abcg2* mRNA expression levels within the whole cohort. We corrected for multiple comparisons using the false discovery rate Benjamini Hochberg method and deemed corrected p-values (q-values) that were less than 0.05 to be significant. Heatmaps were generated to visualise associations between relative bacterial abundance and continuous variables. A hierarchical clustering technique was used to summarise these relationships in the form of a dendogram. Biologically similar communities have a shorter branch length between them.

### Assessment of features that contribute to gut homeostasis and discriminate mothers

We evaluated the cohort using principal components analysis (PCA) based on correlations to determine if we could discriminate mothers based on features related to variation in gut homeostasis: microbial abundance dissimilarity (using mean wUniFrac distance); SI IEL inflammatory load (using a summary score of SI IEL inflammatory biomarkers); relative index of proliferation (Ki67) in SI villi, and MDR mRNA expression (for both *Abcb1a* and *Abcg2* genes). The SI IEL inflammation summary score was calculated by ranking each of the 7 SI IEL inflammatory biomarkers that were significantly different between dietary groups from lowest to highest concentration for all 18 mothers in the cohort for which there were inflammatory measures. Rank values ranged from 1 to 18, where higher values indicate greater inflammatory load. For each mother, a single summary score was calculated by summing the ranked value across each of the 7 biomarkers to derive a single summary score of SI IEL inflammatory load. Loadings of all features (variables) of each principal component (PC) were determined, and loadings most influential for each PC (i.e. the features that contributed the most to the PC) were identified and are displayed in a table of loadings (Figure A.1).

## Results

### Maternal malnutrition alters intestinal architecture and cellular proliferation

Examination of maternal SI gross morphology revealed typical intestine histological components, i.e. the mucosa being formed by regular villi, lamina propria, crypt and goblet cells, and the submucosa formed by its typical inner and outer muscularis layers. In contrast, SI architecture was disrupted by maternal malnutrition. Maternal SI crypt depth was longer in HF mothers compared to CON and UN (p=0.0002, Figure 1), a 10.6% increase in depth vs. CON alone in HF (*post hoc* p=0.02). Crypt depth was shorter (8.6% decrease) in UN mothers compared to CON, but differences were not statistically significant. Villus height and villus:crypt length ratio were not altered by diet (Figure 1). Cellular proliferation in maternal SI crypts and villi were assessed by Ki67 staining (Figure 2A). The mean relative index of proliferation was significantly greater in UN villi (83% ± 7) vs. HF (52% ± 13) and CON (43% ± 15) (p<0.0001, Figure 2B). There were no differences in proliferation index in the crypts across dietary groups (75-79%, Figure 2B).

**Figure 1.**
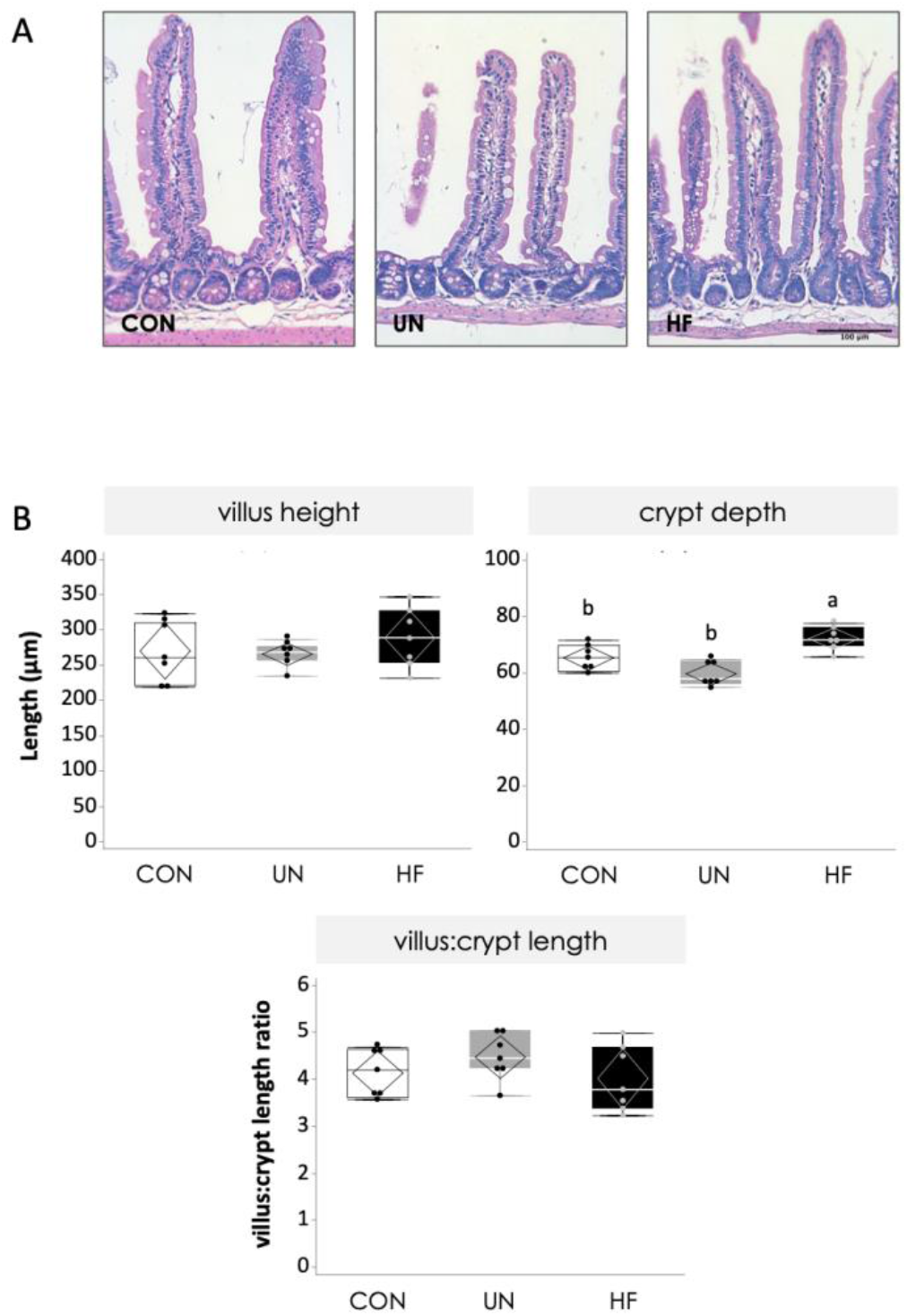
Maternal malnutrition affects small intestinal (SI) architecture and cellular proliferation at GD18.5. A. Representative images of H&E stained maternal SI at 20X magnification. Scale bar = 100 µm. B. Length of villi and crypts (upper panel) and villus:crypt length ratio (lower panel). Data are quantile box plots with 95% CI confidence diamonds.

**Figure 2.**
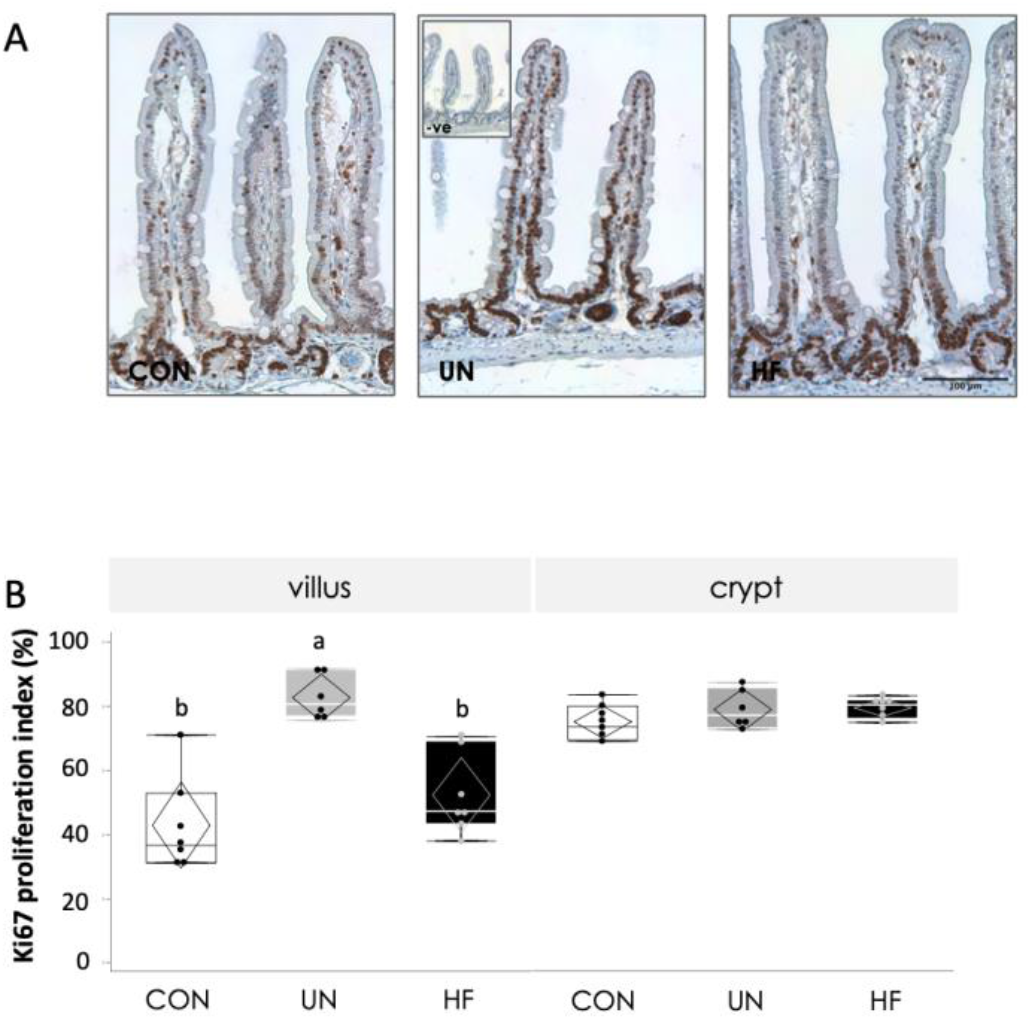
Maternal undernutrition affects cellular proliferation in the small intestine at GD18.5. A. Representative images of immunoreactive Ki67 staining maternal SI at 20X magnification with negative control (inset). Scale bar = 100 µm. B. Proliferation index in villi and crypts. Data are quantile box plots with 95% CI confidence diamonds.

### Maternal HF diet elicits an inflammatory response locally in the SI

To determine whether our models of UN and HF diets were associated with local or systemic inflammation, we measured a panel of pro-inflammatory cytokines and chemokines in maternal SI IEL and plasma. Levels of the pro-inflammatory cytokines and chemokines IL-1α, IL-1β, GM-CSF, IFNγ, eotaxin and MCP-1 were markedly increased in SI IEL of HF compared to CON and UN mothers (p<0.05; Table 1). IL-2 levels were also increased in HF compared to UN mothers (p<0.05, Table 1), and there was an overall difference in levels of the pro-inflammatory cytokines IL-12 (p40) and IL-12 (p70) and the anti-inflammatory cytokine IL-10 between dietary groups (p<0.05), but no between group differences on post hoc analyses (Table 1). A visual synthesis of the significantly different inflammatory biomarker levels in maternal SI IEL clearly showed HF fed mothers had greater inflammatory load compared to CON and UN (Figure 3). Systemic inflammatory load was only modestly altered in a diet-dependent manner. Plasma levels of IL-12 (p40) were reduced in UN mothers and increased in HF mothers (p<0.0001) compared to CON (Table 2). Plasma levels of the chemokine RANTES, a pro-inflammatory mediator, were also increased in HF mothers compared to CON (p<0.02, Table 2).

**Table 1.** Inflammatory biomarkers in maternal small intestinal intraepithelial lymphocytes at GD18.5.

| Biomarker (pg/mg protein) | CON | UN | HF | p-value |
| --- | --- | --- | --- | --- |
| IL-1a | 2.8 (1.9-5.2)<sup>b</sup> | 2.5 (2.2-4.7)<sup>b</sup> | 10.0 (5.6-16)<sup>a</sup> | 0.009 |
| IL-1b | 156 ± 103<sup>b</sup> | 144 ± 44<sup>b</sup> | 288 ± 93<sup>a</sup> | 0.02 |
| IL-2 | 6.6 (5.3-14)<sup>ab</sup> | 6.4 (4.9-12)<sup>b</sup> | 16.6 (11-19)<sup>a</sup> | 0.03 |
| IL-6 | 3.8 (1.9-7.3) | 7.3 (3.3-11) | 3.5 (1.9-10) | NS |
| IL-10 | 4.3 ± 2.4 | 4.1 ± 1.9 | 7.4 ± 2.7 | 0.049 |
| IL-12 (p40) | 1.8 (1.4-2.7) | 2.0 (1.7-2.2) | 3.0 (2.6-4.5) | 0.044 |
| IL-12 (p70) | 82.6 ± 64 | 72.5 ± 37 | 141 ± 37 | 0.046 |
| Eotaxin | 459 ± 197<sup>b</sup> | 447 ± 109<sup>b</sup> | 741 ± 211<sup>a</sup> | 0.02 |
| GM-CSF | 66.7 ± 32<sup>b</sup> | 65.9 ± 13<sup>b</sup> | 130 ± 37<sup>a</sup> | 0.002 |
| IFN-γ | 4.0 (3.3-5.9)<sup>b</sup> | 4.2 (3.5-5.1)<sup>b</sup> | 8.3 (7.6-14)<sup>a</sup> | 0.0006 |
| MCP-1 | 6.4 (4.9-9.9)<sup>b</sup> | 6.4 (5.7-10)<sup>b</sup> | 12.2 (10-17)<sup>a</sup> | 0.01 |
| MIP-1a | 14.5 ± 9.9 | 13.3 ± 4.5 | 20.3 ± 6.5 | NS |
| MIP-1b | 3.5 ± 2.1 | 4.1 ± 0.72 | 4.4 ± 1.9 | NS |
| RANTES | 7.2 (5.6-11) | 6.7 (4.4-7.7) | 6.4 (3.4-13) | NS |
| TNF-a | 49.0 (27-189) | 126 (65-193) | 67.0 (42-192) | NS |

Data are mean ± SD or median (IQR). Groups with different letters are significantly different (Tukey's *post hoc*).

**Table 2.** Inflammatory biomarkers in maternal plasma at GD18.5.

| Biomarker (pg/mL) | CON | UN | HF | p-value |
| --- | --- | --- | --- | --- |
| IL-1a | 37.0 ± 21 | 40.5 ± 28 | 37.7 ± 20 | NS |
| IL-1b | 199 (73-344) | 251 (75-364) | 330 (206-426) | NS |
| IL-2 | 55.5 ± 32 | 64.8 ± 33 | 61.8 ± 17 | NS |
| IL-5 | 20.5 ± 5.7 | 17.1 ± 13 | 23.8 ± 11 | NS |
| IL-6 | 10.5 ± 7.1 | 10.2 ± 6.1 | 12.1 ± 4.3 | NS |
| IL-10 | 52.4 ± 48 | 76.3 ± 67 | 100 ± 41 | NS |
| IL-12 (p40) | 138 ± 55<sup>a</sup> | 73.8 ± 42<sup>b</sup> | 207 ± 27<sup>c</sup> | <0.0001 |
| IL-12 (p70) | 105 ± 133 | 218 ± 236 | 271 ± 128 | NS |
| IL-17 | 88.1 ± 69 | 137 ± 100 | 151 ± 52 | NS |
| Eotaxin | 941 ± 789 | 1449 ± 774 | 1048 ± 673 | NS |
| G-CSF | 156 (62-172) | 120 (56-170) | 131 (128-165) | NS |
| GM-CSF | 280 ± 112 | 239 ± 102 | 251 ± 120 | NS |
| IFN-γ | 20.2 ± 16 | 29.8 ± 33 | 23.7 ± 15 | NS |
| KC | 25.5 ± 7.9 | 25.0 ± 6.2 | 28.2 ± 12 | NS |
| MCP-1 | 88.8 ± 72 | 120 ± 88 | 143 ± 63 | NS |
| MIP-1a | 45.0 ± 13 | 43.7 ± 37 | 53.7 ± 15 | NS |
| MIP-1b | 32.9 ± 11 | 29.8 ± 20 | 32.5 ± 10 | NS |
| RANTES | 13.5 ± 5.0<sup>a</sup> | 14.3 ± 11<sup>ab</sup> | 22.8 ± 1.4<sup>b</sup> | 0.02 |
| TNF-a | 489 ± 480 | 470 ± 706 | 816 ± 361 | NS |

Data are mean ± SD or median (IQR). Groups with different letters are significantly different (Tukey's or Games-Howell *post hoc*).

**Figure 3.**
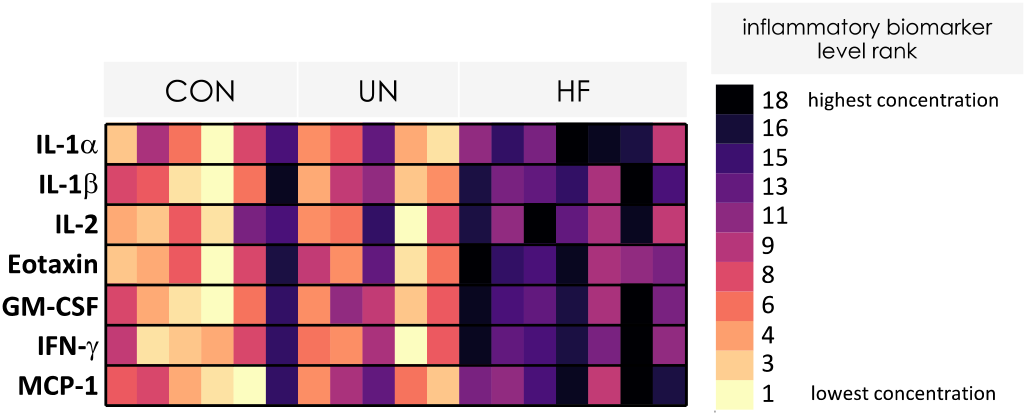
Heatmap of inflammatory biomarker level rank for each of the seven SI IEL biomarkers that were significantly different between dietary groups (refer to Table 1). For each inflammatory biomarker, concentrations were ranked from lowest to highest concentration for all mothers. Rank values ranged from 1 to 18, where higher values indicate higher concentrations for that inflammatory biomarker in SI IEL.

### Maternal malnutrition alters gut mRNA expression of multidrug resistance transporters

To determine whether maternal malnutrition can alter P-gp and BCRP transport potential in the gut, we measured the expression levels and determined the maternal SI localisation of the multidrug resistance transporters P-gp/*Abcb1a/b* and BCRP/*Abcg2*. UN was associated with a 3.7-fold increase in *Abcb1a* mRNA expression in maternal SI compared to CON and HF (p=0.0002, Figure 4A) and a decrease in *Abcg2* mRNA expression in maternal SI compared to CON and HF (p=0.004, Figure 4C). There was no effect of maternal UN on *Abcb1b* mRNA expression in maternal SI (Figure 4B). In the fetal gut, we did not observe any differences in *Abcb1a/b* or *Abcg2* expression levels due to maternal UN or HF diets when analysing all fetuses, inclusive of sex (Figure 4), or when stratifying by fetal sex (Table A.2). Additionally, to determine whether the effects of maternal diet were tissue specific, we evaluated mRNA expression of our genes of interest in the maternal cerebella. HF dams showed reduced cerebellar *Abcb1b* mRNA expression compared to CON (p=0.04. Figure 4B) and there was an overall slight difference in *Abcg2* mRNA expression in cerebella (p=0.046), but there were no significant differences between dietary groups on post hoc testing (Figure 4C). To establish whether alterations in maternal diet were related to changes in the distribution of P-gp and BCRP protein within the maternal intestinal mucosa (crypt and villi) and within the epithelial cell (membranous versus cytoplasmic), immunohistochemical analysis was performed. Immunoreactive (ir)-P-gp signal was detected in the villi and crypt epithelia in all mothers, with strongest staining localised to the apical membrane of the intestinal villi and weaker staining in the cytoplasm (Figure 5A-C). ir-BCRP signal was present in villi and crypt surface epithelium and cytoplasm (Figure 5D-F). The cellular localisation of P-gp and BCRP proteins along the villi and crypts was similar in all mothers, irrespective of diet (Figure 5A-C).

**Figure 4.**
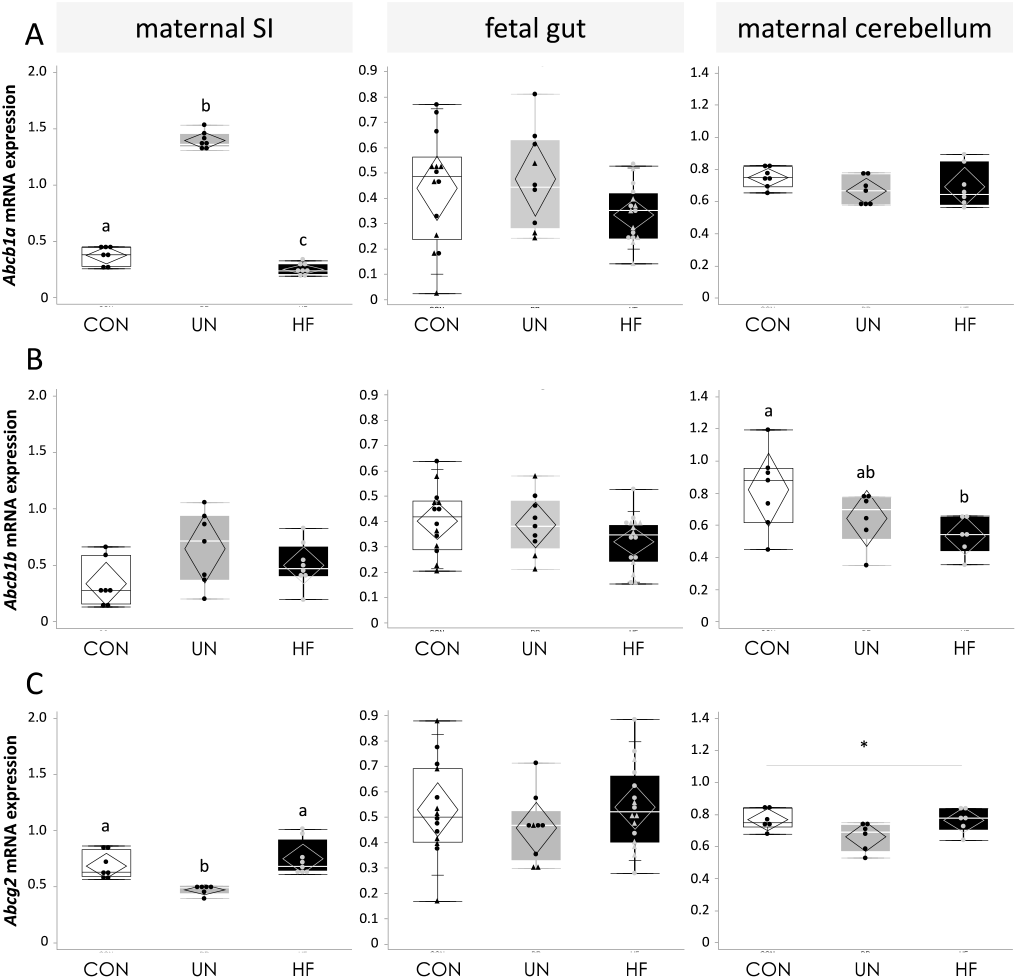
MDR mRNA expression in CON, UN, and HF dams and fetuses at GD18.5. A. *Abcb1a* mRNA expression levels in maternal SI, fetal gut, and maternal cerebellum. B. *Abcb1* mRNA expression levels in maternal SI, fetal gut, and maternal cerebellum. C. *Abcg2* mRNA expression levels in maternal SI, fetal gut, and maternal cerebellum. Data are quantile box plots with 95% CI confidence diamonds. *p<0.05 (ANOVA). Groups with different letters are significantly different (Games-Howell or Steel-Dwass *post hoc*).

**Figure 5.**
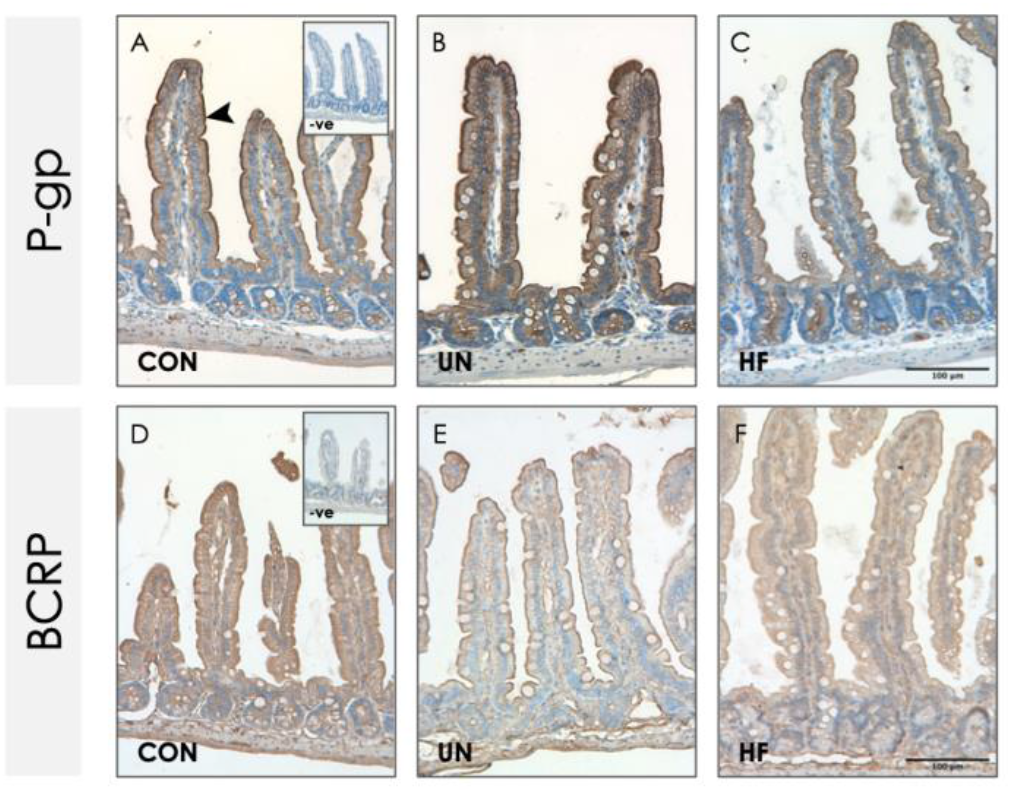
Small intestinal MDR protein expression in CON, UN, and HF dams at GD18.5. Representative images of immunoreactive P-gp (A-C) and BCRP (D-F) staining maternal SI with negative controls (inset). Images captured at 20X magnification. Scale bar = 100 µm. Large arrowhead = apical membrane.

### Expression of Abcb1a and Abcg2 correlates with SI villus proliferation index in UN pregnancies

As we observed the greatest change in the relative proliferation index and MDR transporter mRNA levels in the SI of UN pregnancies, we conducted a correlation analysis between these variables and observed that villus proliferative status was associated with drug transport expression. There was a significant positive relationship between the relative index of proliferation in the villus and *Abcb1a* mRNA expression (rho=0.53, p=0.017, Figure 6A), but not in the crypt (rho=0.02; p=NS). In contrast, there was a significant negative relationship between the relative index of proliferation in the villus and *Abcg2* mRNA expression (rho=-0.65, p=0.002, Figure 6B), but not in the crypt (rho=-0.07, p=NS).

**Figure 6.**
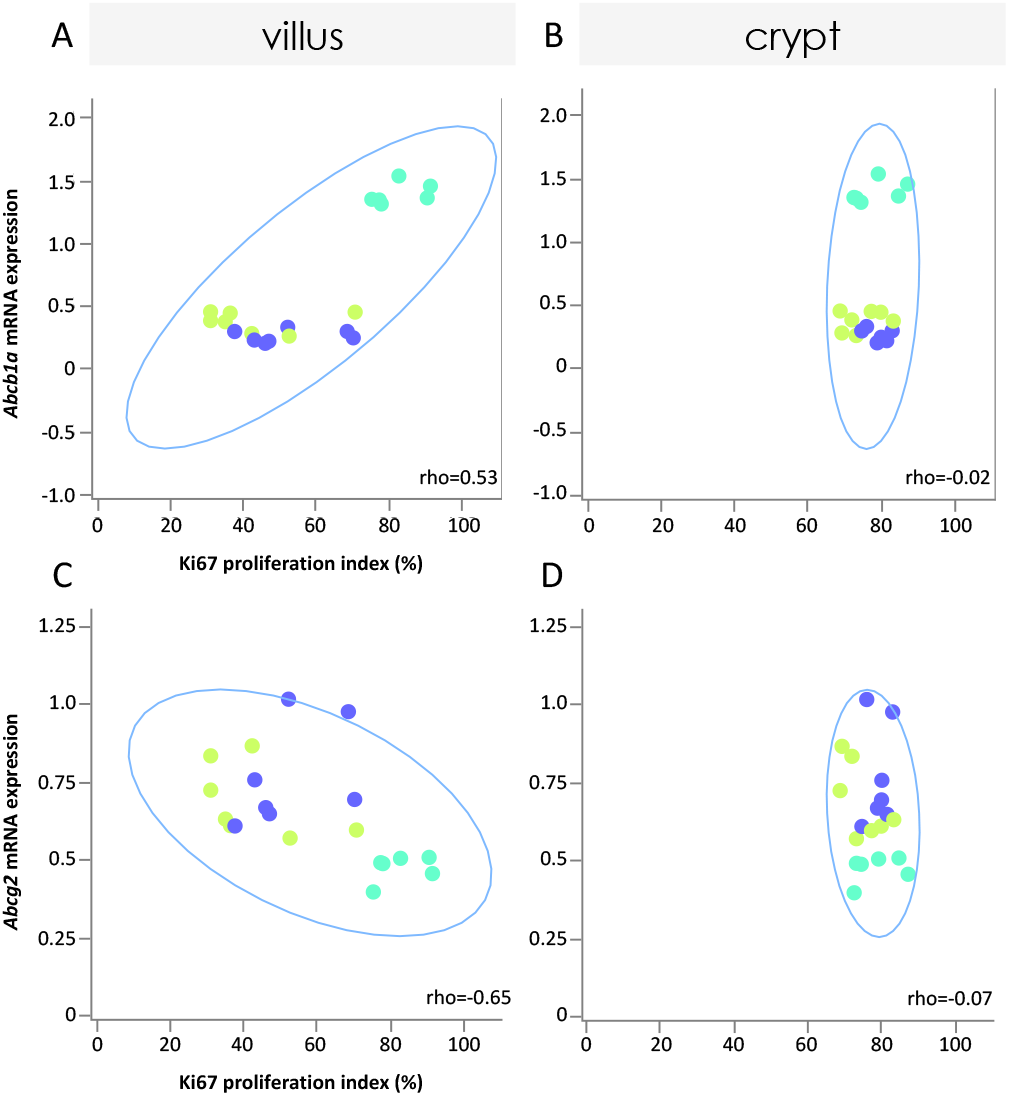
Relationships between relative index of proliferation in maternal villi and crypts with *Abcb1a* (A, B) and *Abcg2* (C, D) mRNA expression levels, inclusive of all diets. Lime-yellow circles = CON, blue circles = HF, teal circles = UN. 95% density ellipses shown in light blue. p<0.05 (Spearman’s rho).

### Maternal malnutrition alters the gut microenvironment, and these changes are associated with drug transporter expression

Since we previously showed that maternal malnutrition, particularly HF diet, alters the composition of the gut microbiome[35], and herein we observed that maternal malnutrition modulates gut mRNA expression of key MDR transporters, we sought to determine if there were relationships between maternal gut microbiome composition and MDR transporter expression. Inclusive of all diets, we found significant associations between *Abcb1a* and *Abcg2* mRNA expression levels with relative abundance of microbial taxa, where the direction of relationship was transporter-dependent. Seven families were significantly positively associated, and seven were significantly negatively associated, with *Abcb1a* mRNA expression levels (q<0.05; Figure 7A). Three families were significantly positively associated, and six families were significantly negatively associated, with *Abcg2* mRNA expression levels (q<0.05; Figure 7A). At the genus level, four genera were significantly positively associated, and six were significantly negatively associated, with *Abcb1a* mRNA expression levels (q<0.05; Figure 7B). Two genera each were significantly positively and negatively associated with *Abcg2* mRNA expression levels (q<0.05; Figure 7B). The relative abundance level of one species, *Clostridium septicum*, was negatively associated with SI *Abcb1a* mRNA expression levels (rho=-0.81; q<0.0004; Figure 8A). Additionally, *C. septicum* relative abundance levels were significantly reduced in UN mothers compared to CON and HF (p=0.006; Figure 8B). There were no significant associations between *Abcg2* mRNA expression levels and species relative abundance levels within the whole cohort. However, diet-stratified analyses identified one classified species, *Trichodesmium erythraeum*, whose relative abundance level was positively associated with SI *Abcg2* mRNA expression levels in HF mothers (rho=1; q=0.005; Figure 8A shows relationship in all mothers). Moreover, relative abundance levels of this species were significantly reduced in HF-fed mothers compared to CON and UN (p=0.004; Figure 8B).

**Figure 7.**
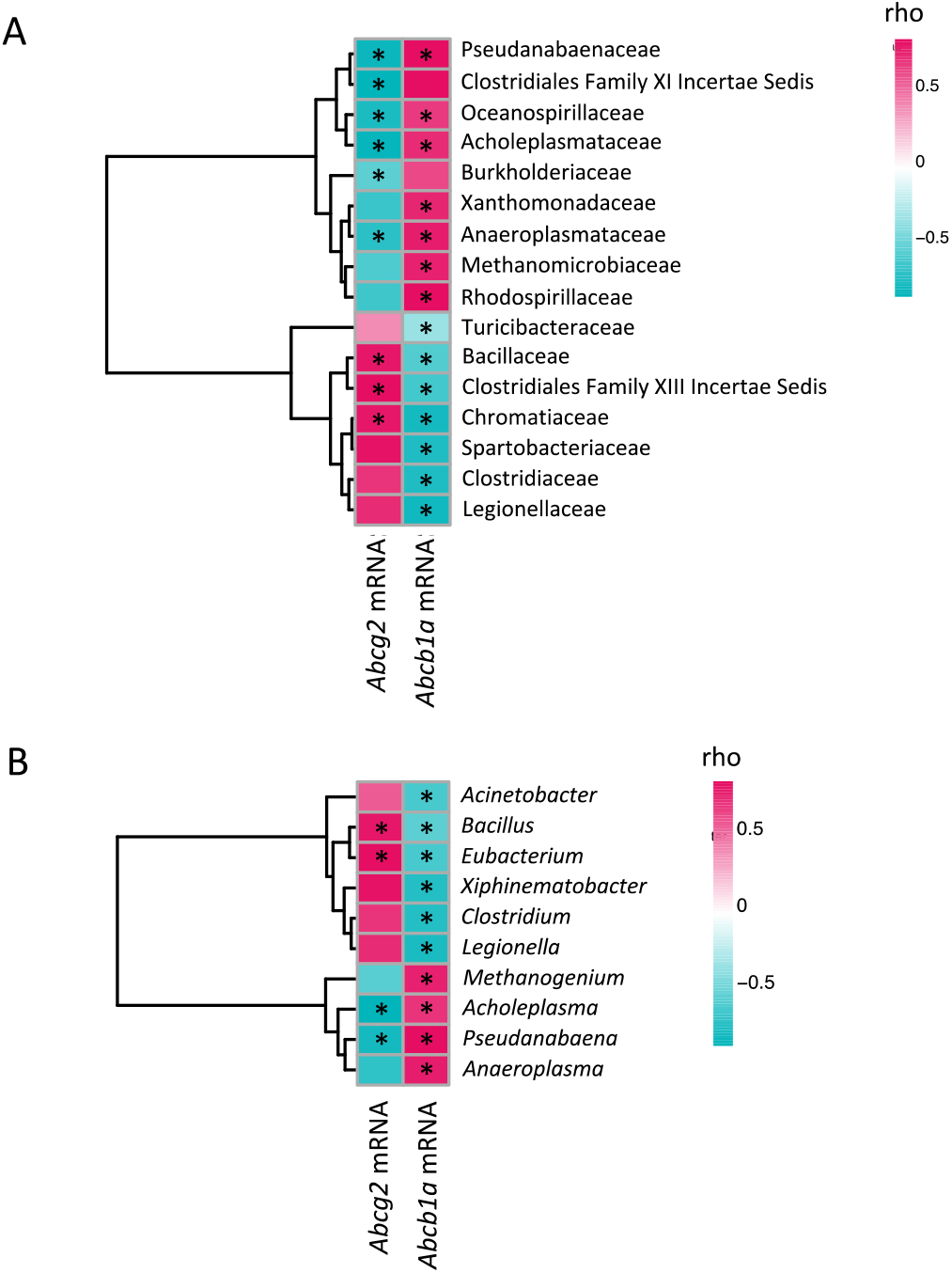
Associations between maternal gut microbial taxa relative abundance at the family (A) and genus (B) levels with efflux transporter mRNA expression levels. Red indicates a positive association, blue indicates a negative association. Colour saturation indicates the strength of relationship. * indicates significant association (q<0.05). See Tables A.3 and A.4 for details.

**Figure 8.**
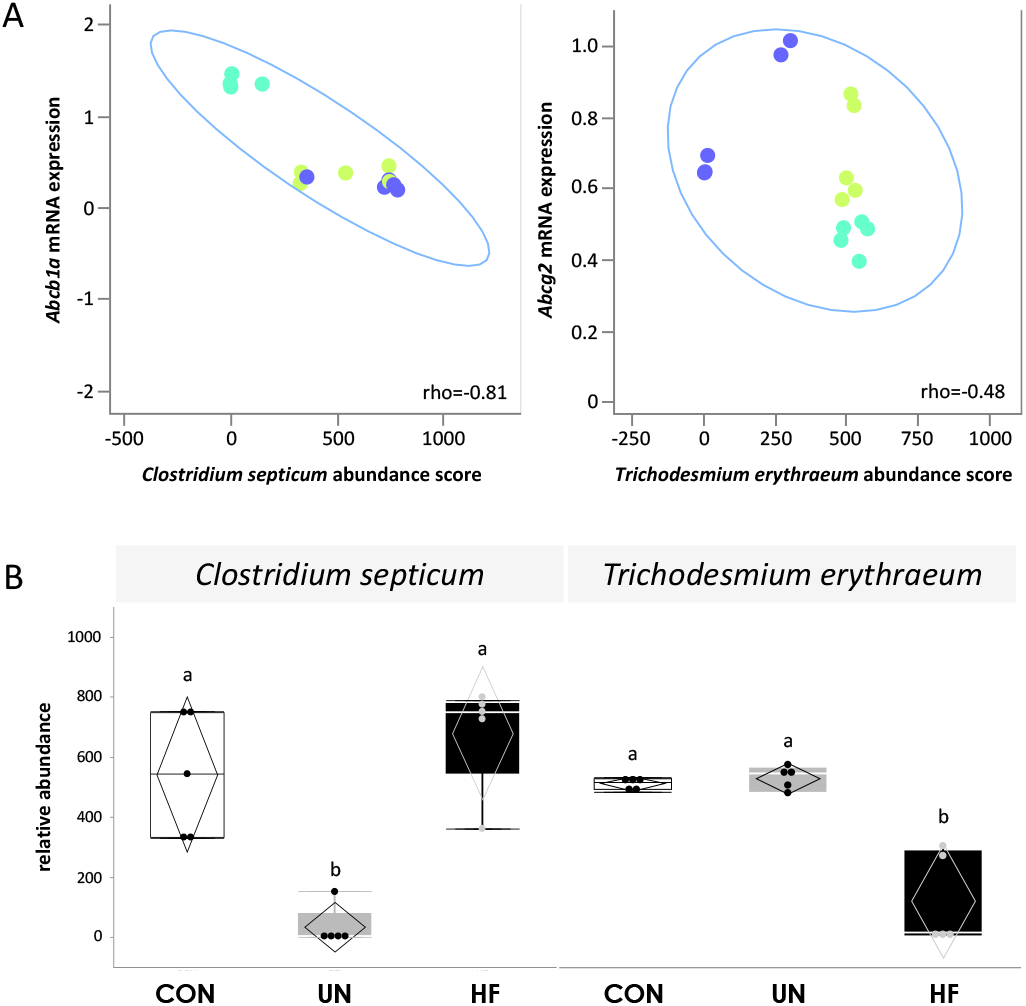
Bacterial species associated with SI MDR mRNA expression. A. Relationships between species relative abundance and *Abcb1a* or *Abcg2* mRNA expression levels in all mothers. Lime-yellow circles = CON, blue circles = HF, teal circles = UN. 95% density ellipses shown in light blue. q<0.05 (Spearman’s rho). *T. erythraeum* relative abundance was positively associated with *Abcg2* mRNA levels in HF mothers (q=0.005). B. Relative abundance levels of *C. septicum* and T. *erythraeum* in maternal caecal contents at GD18.5. Data are quantile box plots with 95% CI confidence diamonds. *p<0.05 (ANOVA). Groups with different letters are significantly different (Steel-Dwass or Games-Howell *post hoc*).

### Maternal diet coordinates the gut holobiome and microenvironment

Principal component analysis revealed that mothers could be discriminated based on their nutritional history when assessing key features contributing to variation in gut homeostasis (gut microbiome dissimilarity, small intestinal inflammatory load and proliferation, and expression levels of MDR transporters). Distinct clusters of mothers emerged primarily based on PC1 (Figure 9). All features were considered important for PC1 (eigenvalue [EV] 2.69, explaining 53.8% of variance in the data, Figure 9 and Figure A.1). Based on PC2, the features with the greatest EVs were represented by gut microbiome dissimilarity, SI IEL inflammatory load, and Ki67 proliferation index in villi (EV 1.24, explaining 24.9% of variance in the data, Figure 9 and Figure A.1). Collectively, PCA revealed that mothers with similar holobionts and gut microenvironments clustered together, and clustering was largely determined by nutritional history.

**Figure 9.**
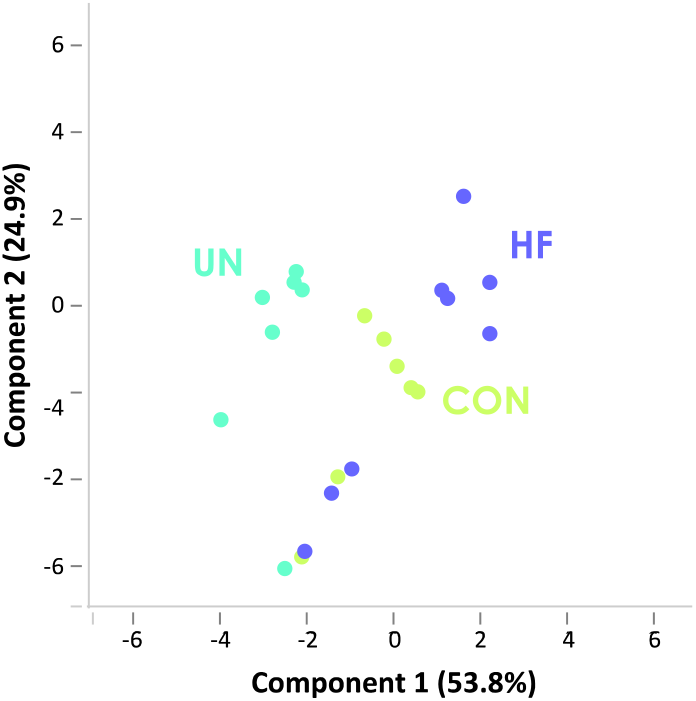
Maternal diet coordinates gut allostasis. PCA of features contributing to maternal gut allostasis. Percent variation explained by the principle components is indicated in parentheses on the x- and y-axes.

## Discussion

Here, we report that maternal diet provokes adaptations in the gut holobiont during pregnancy, improving our understanding of how common nutritional adversities impact maternal gut homeostasis and the mechanisms that contribute to reprogramming the normal course of fetal development. Using a systems physiology approach we identified that malnutrition alters SI architecture and villi proliferation, pro-inflammatory load, and expression of *Abcb1a* and *Abcg2*, key MDR transport systems in pregnancy, without altering their protein localisation. We also report significant associations between *Abcb1a* and *Abcg2* mRNA levels with relative abundance of specific microbial taxa, and that features of the gut microenvironment may be influenced by maternal diet.

We provide the first report of diet-induced changes in MDR transporters in the maternal SI during pregnancy, and their relationships with the gut microbiome. Whilst it is well known that physiologic changes due to pregnancy[48], and malnutrition[24] or metabolic disease[25, 26] in the non-pregnant state affect drug disposition, to date, few studies have explored how maternal gut MDR transport adapts to dietary challenges that also influence pregnancy phenotype. Consistent with our mRNA and immunoreactivity findings, a study in male mice using a similar degree of caloric restriction as reported here, found increased *Abcb1a* mRNA expression in the duodenum and jejunum of calorically-restricted mice, with the most striking increase in expression in the jejunum[24]. P-gp activity was also shown to be increased in these mice[24]. In contrast, *Abcb1a* mRNA expression was lower in HF mothers in our study, with levels consistent with lower *Abcb1a* expression we have reported in HF placentae[3]. We also report lower *Abcg2* mRNA expression in SI of UN mothers, consistent with changes in placentae from UN mothers[3] and from pregnancies complicated by intrauterine growth restriction[49]. Conversely, we did not observe any changes in *Abcg2* expression in SI of HF mothers, despite that BCRP expression has been reported to be lower in the SI mucosa of HF-fed male mice[50], highlighting that pregnancy may impact *Abcg2* SI responsiveness to maternal malnutrition. Importantly, studies in rodents have shown that site-specific mRNA expression levels of P-gp and BCRP are good correlates with protein expression[19]. P-gp substrates include antibiotics (amoxicillin, clarithromycin, ciprofloxacin, levofloxacin, rifampicin, tetracycline), antiretrovirals (indinavir, saquinavir, ritonavir, nelfinavir), antifungals (itraconazole), stomach-protective drugs (cimetidine, ranitidine), nonsteroidal anti-inflammatory drugs (NSAIDs; diclofenac), synthetic glucocorticoids (dexamethasone prednisolone), cytokines (IL-6; IFN-γ; TNF-α) and nutrients (flavonoids), and BCRP substrates include antibiotics (beta-lactams, fluoroquinolones), antiretrovirals (indinavir, zidovudine), sulfonylureas (glyburide) and nutrients (folate)[22]. Modulation of these transporters in the maternal gut by diet can potentially alter the local and systemic levels of their substrates. Thus, our findings suggest there may be implications not only for the availability and biodisposition of drugs, but also nutrients and inflammatory molecules, and thus potential fetal exposure to these, if the mother is malnourished. Future studies should investigate the effect of pregnancy alone and with dietary challenge (vs. the non-pregnant basal and challenged states in females) in modulating MDR transporter systems in enterocytes and their relationship with maternal physiology.

The cellular and molecular mechanisms explaining how malnutrition alters MDR expression in pregnancy are unclear. There is limited information demonstrating direct regulation of MDR transporters by nutrients. However, MDRs do interact with a number of substrates which may be derived directly from the diet, and or/dietary components may influence the production/levels of substrates and other molecules that interact directly with MDRs, as seen with inflammation (and thus, metainflammation)[22], and cell cycle stage and p53 expression[51, 52]. To address this, we first explored whether and how maternal gut morphology adapts to the spectrum of malnutrition, as intestinal architecture and morphology may influence cellular function. We observed a greater crypt depth in SI from HF mothers compared to CON. Crypt hyperplasia, hallmarked by an elongated crypt due to an increase in the number of epithelial cells, is one defining feature of intestinal inflammation[53]. Whilst we did not observe an increase in the proliferative index of HF SI compared to CON, we did show an increase in the levels of seven pro-inflammatory biomarkers in SI IEL, specific to HF pregnancies. Intestinal IELs are considered to be gut sentinels, which effectively contribute to the maintenance of mucosal barrier integrity in part through their production of a number of cytokines that regulate intestinal barrier homeostasis[54]. When dysregulated, IELs may promote the onset of gut and systemic pathologies[55]. In *Mdr1a* ^*-/-*^ mice, HF diet worsens spontaneous inflammatory bowel disease and histologic pathologies, including elongation of crypts[39]. Notably, *ABCB1*/P-gp expression levels are lower in inflamed intestinal epithelia of patients with gastrointestinal diseases (vs. uninflamed intestinal epithelia), and *in vitro*, lower *ABCB1* expression has been observed with higher levels of cytokines[56]. Thus, the pro-inflammatory environment established by HF feeding may contribute to a downregulation of *Abcba1* expression in our model. This would be consistent with the observed altered expression or activity of these MDRs in the yolk sac, placenta and fetal blood brain barrier in the presence of bacterial and viral infection[57–59] including malaria[60], resulting in increased fetoplacental substrate accumulation, altered fetal growth, or preterm delivery.

In contrast to the HF diet, the villus appeared impacted by maternal UN. We observed increased proliferation in UN SI, consistent with other caloric restriction studies in the non-pregnant host[61]. Increased proliferation in villi is an adaptation aimed to improve the absorptive capacity of the intestine, particularly in an effort to sequester more nutrients when bodyweight decreases[61]. We determined that there was a positive relationship between villus proliferation index and *Abcb1a* mRNA expression, largely driven by maternal UN, and a negative relationship between villus proliferation index and *Abcg2* mRNA expression. These findings are consistent with studies that show targeted silencing of *ABCB1* results in reduced cell proliferation and invasion in human extravillous trophoblast (EVT)[62], and BCRP inhibits EVT cell migration[63], whilst P-gp overexpression in fibroblasts accelerates cell proliferation and migration[64]. Therefore, MDR expression may play a key role not only in substrate transport dynamics, but also maintenance of key barrier tissues, especially those that are rapidly developing or turning over. Altered cellular proliferation and migration may indirectly alter the absorption and excretion of important pharmacological and physiological substrates during reproduction and pregnancy. Additionally, of importance, we did not observe any changes in fetal gut mRNA expression of these MDRs. This, in consideration with our previous findings of altered placental *Abcb1a* and *Abcg2* mRNA expression in placentae from malnourished mothers[3], suggests that fetal gut MDR is not programmed by maternal diet in term pregnancies, although altered maternal gut and placental MDR expression and function could contribute to fetal maldevelopment independent of changes in fetal gut MDRs, through changes in transport and bioavailability of P-gp and BCRP substrates.

To further understand possible mechanisms underlying the observed changes in intestinal allostasis, we next explored whether there were relationships between maternal SI MDR expression levels and gut bacteria. MDRs are expressed across the gastrointestinal tract, including in the jejunum[65, 66] and are strategically localised[19–21] (at the enterocyte apical membrane) to play key roles in host-microbe interactions in the gut, although beyond pathogenic bacteria[67, 68], whether there is a functional consequence of microbial-MDR cross talk is not known. Here we provide the first report of relationships between SI MDR transporters and microbial relative abundance levels in malnourished pregnancies. We found significant associations between relative abundance levels of specific taxa and expression of *Abcb1a* and *Abcg2*, including a negative association between *C. septicum*, a Gram-positive spore-forming bacillus in the human and animal GI tract[69], and *Abcb1a*. When exploring these relationships more closely relative to maternal diet, we found the relative abundance of this species was lower in UN mothers compared to CON and HF. Members of the Clostridia class have key roles in gut homeostasis and maintenance of mucous production by the host[70–72], and have recently been shown to correlate with colonic P-gp expression in mice[73]. We previously found increased goblet cell number in UN maternal SI[34], which collectively with higher *Abcb1a* expression, may suggest a beneficial adaptation of the UN gut to increase intestinal defences, including to potentially harmful microbes or their metabolites. In contrast, *Abcg2* expression was positively associated with relative abundance levels of *T. erythraeum*, but in HF mothers only, where abundance levels were significantly reduced compared to CON and UN. We previously reported relationships between *T. erythraeum* relative abundance levels and concentrations of maternal circulating inflammatory and metabolic biomarkers[35], and noted that although little is known about this species in the context of mammalian health, it is important for iron storage[74] and DNA protection against oxidative stress[75, 76]. BCRP/*Abcg2* can efflux porphyrin/haem in an effort to tightly regulate iron availability for host cellular metabolism and prevent toxicity[77, 78]. Abundance levels of microbes will be influenced by iron availability, in part determined by mechanisms that regulate host iron biodisposition, including efflux transport. It is also plausible that gut microbes or their metabolites could directly influence these mechanisms[79] [73], and the direct regulation of intestinal MDRs by microbes should be evaluated in pregnancy under basal and challenged conditions, including poor nutrition. Additionally, future studies should test whether any of the inflammatory, histomorphological, functional and/or microbial changes in the gut of HF-fed mothers can be corrected by transplanting microbes from CON or UN mothers, similar to studies showing a beneficial effect of diet and exercise on host phenotype following fecal transplantation[80]. In doing so, we may not only identify a target for the prevention of these outcomes, but also better understand the mechanisms that permit specific holobiont maladaptations.

A limitation of our study is that we did not measure the SI microbiome, but suggest that the caecal microbiome can inform about SI function. There is limited understanding of the human and mouse SI microbiomes, including how microbes residing in the SI may influence key physiological functions in their niche, such as nutrient metabolism or epithelial defences. Mice are coprophagic, and a series of elegant studies showed that this behaviour alters the SI microbial load and composition and function of the SI microbiome, with limited or no effects in the lower gastrointestinal tract[81]. Further, coprophagic mice have SI and stomach microbiomes that are compositionally more similar to caecal microbiomes than non-coprophagic mice[81]. Studies have shown that the relative abundance of major phyla in the mouse SI, colon and caecum are similar, but there are genera exclusive to each niche[82, 83]. We believe sampling the caecal contents can provide a reasonable picture of gut microbial changes to environmental conditions[84], including diet, that cannot easily be ascertained from limited samples in the upper gastrointestinal tract. Future studies should consider the role of coprophagy and the SI microbiome on host response to malnutrition[85]. Another potential limitation is the nature of P-gp and BCRP gut protein immunohistochemistry evaluations, which were undertaken to assess whether maternal malnutrition during pregnancy could alter localisation of these proteins. We did not observe alterations in P-gp and BCRP localisation among groups. Further, we did not use Western blot to quantify protein changes of P-gp and BCRP as this approach would reflect the overall protein levels in whole cells (including cytoplasm, organelles and nucleus) and would potentially mask differences in plasma membrane protein levels between experimental groups. Rather, we used immunohistochemistry to assess P-gp and BCRP staining in the apical (luminal-facing) membrane of enterocytes. Future studies should investigate how maternal malnutrition impacts P-gp and BCRP expression in isolated apical membranes of enterocytes, and evaluate enterocyte P-gp and BCRP function in response to dietary challenges and changes in the holobiont during pregnancy.

Our study identifies adaptations in the pregnancy gut holobiont to dietary challenge that could affect pregnancy health and contribute to fetal maldevelopment. We suggest attempts to improve outcomes for pregnancies and offspring exposed to adverse dietary conditions have only partially been successful due to the complexity of interactions between the environment, host, and its resident microbes. Here we took a systems physiology approach to better understand these relationships, and propose that gut holobiont dysfunction and diet—MDR—microbe interactions may be key targets for improving reproductive and pregnancy health, particularly in mothers with an already imbalanced gut holobiont.

## Acknowledgments

The authors wish to thank Ela Matysiak and Richard Maganga for their assistance with the animal cohort and Adam Altrichter for his assistance with the microbiome analyses.

## Author Contributions

Conceptualisation, KLC, SJL, EB; data acquisition/analysis KLC, EB, TZD; writing/revising KLC, EB, SJL, TZD.

## Supplementary files

This article contains supplemental figures and tables, which can be found at the end of this document.

---

**Table A.1.**
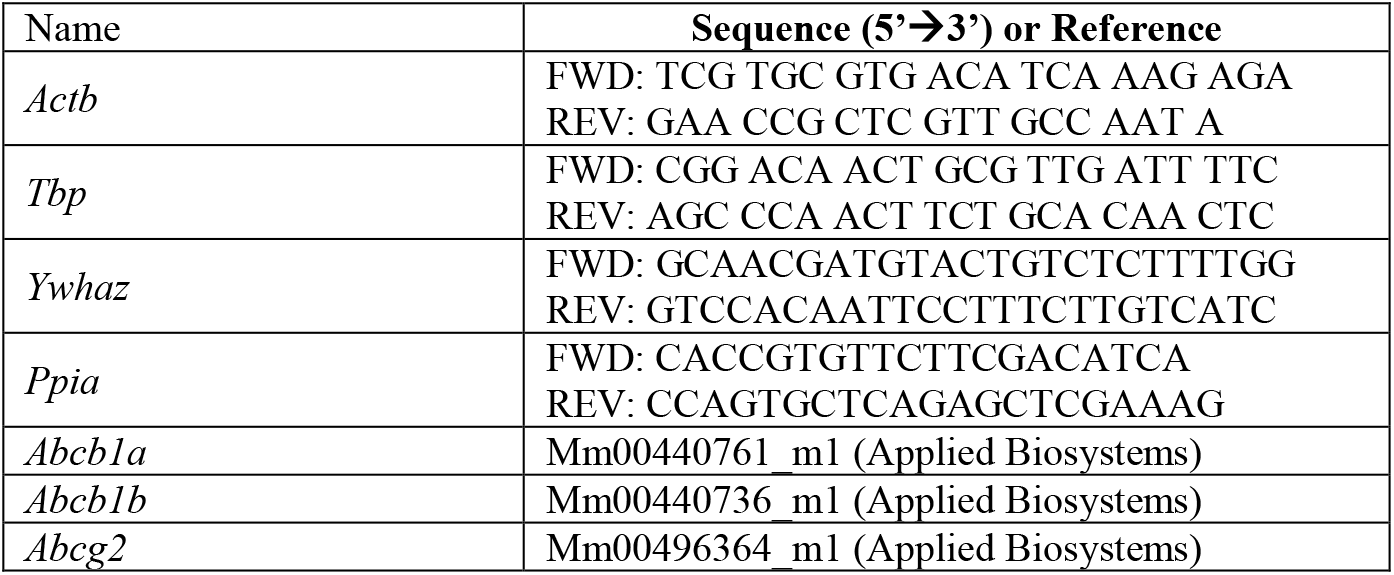
Primer sequences for qPCR

**Table A.2.** Multidrug resistance gene mRNA expression in female and male guts at GD18.5.

|  | Females |  |  |  | Males |  |  |  |
| --- | --- | --- | --- | --- | --- | --- | --- | --- |
|  | CON | UN | HF | p-value | CON | UN | HF | p-value |
| **MDR transporters** |  |  |  |  |  |  |  |  |
| *Abcb1a* mRNA | 0.36 ± 0.20 | 0.35 ± 0.16 | 0.31 ± 0.12 | NS | 0.52 ± 0.22 | 0.54 ± 0.18 | 0.36 ± 0.11 | NS |
| *Abcb1b* mRNA | 0.37 ± 0.14 | 0.35 ± 0.20 | 0.31 ± 0.10 | NS | 0.43 ± 0.11 | 0.40 ± 0.07 | 0.33 ± 0.11 | NS |
| *Abcg2* mRNA | 0.51 ± 0.23 | 0.36 ± 0.10 | 0.48 ± 0.12 | NS | 0.55 ± 0.14 | 0.51 ± 0.12 | 0.60 ± 0.19 | NS |

Data are mean ± SD. NS = not significant.

**Table A.3.** Correlations between microbial family relative abundance levels and ABC transporter mRNA expression.

| Family | Transporter | Correlation coefficient | p value | Corrected q value |
| --- | --- | --- | --- | --- |
| Pseudanabaenaceae | *Abcb1a* | 0.80 | 0.001 | **0.022** |
|  | *Abcg2* | -0.85 | 0.002 | **0.022** |
| Clostridiales Family XI Incertae Sedis | *Abcb1a* | 0.79 | 0.03 | 0.095 |
|  | *Abcg2* | -0.90 | 0.002 | **0.022** |
| Oceanospirillaceae | *Abcb1a* | 0.64 | 0.005 | **0.031** |
|  | *Abcg2* | -0.80 | 0.002 | **0.022** |
| Acholeplasmataceae | *Abcb1a* | 0.69 | 0.003 | **0.023** |
|  | *Abcg2* | -0.87 | 0.0008 | **0.022** |
| Burkholderiaceae | *Abcb1a* | 0.59 | 0.03 | 0.096 |
|  | *Abcg2* | -0.59 | 0.006 | **0.037** |
| Xanthomonadaceae | *Abcb1a* | 0.70 | 0.007 | **0.037** |
|  | *Abcg2* | -0.71 | 0.07 | 0.14 |
| Anaeroplasmataceae | *Abcb1a* | 0.74 | 0.003 | **0.023** |
|  | *Abcg2* | -0.75 | 0.008 | **0.046** |
| Methanomicrobiaceae | *Abcb1a* | 0.73 | 0.005 | **0.034** |
|  | *Abcg2* | -0.63 | 0.07 | 0.15 |
| Rhodospirillaceae | *Abcb1a* | 0.79 | 0.002 | **0.022** |
|  | *Abcg2* | -0.69 | 0.06 | 0.14 |
| Turicibacteraceae | *Abcb1a* | -0.38 | 0.005 | **0.031** |
|  | *Abcg2* | 0.36 | 0.26 | 0.37 |
| Bacillaceae | *Abcb1a* | -0.61 | <0.0001 | **0.0003** |
|  | *Abcg2* | 0.75 | 0.002 | **0.022** |
| Clostridiales Family XIII Incertae Sedis | *Abcb1a* | -0.67 | 0.002 | **0.022** |
|  | *Abcg2* | 0.79 | 0.002 | **0.022** |
| Chromatiaceae | *Abcb1a* | -0.82 | <0.0001 | **0.0003** |
|  | *Abcg2* | 0.77 | 0.004 | **0.031** |
| Spartobacteriaceae | *Abcb1a* | -0.79 | 0.002 | **0.022** |
|  | *Abcg2* | 0.78 | 0.01 | 0.051 |
| Clostridiaceae | *Abcb1a* | -0.78 | 0.0003 | **0.01** |
|  | *Abcg2* | 0.65 | 0.05 | 0.13 |
| Legionellaceae | *Abcb1a* | -0.83 | 0.002 | **0.022** |
|  | *Abcg2* | 0.69 | 0.11 | 0.20 |

Refer to heatmap Figure 7A.

**Table A.4.** Correlations between microbial genus relative abundance levels and ABC transporter mRNA expression.

| Genus | Transporter | Correlation coefficient | p value | Corrected q value |
| --- | --- | --- | --- | --- |
| *Acinetobacter* | *Abcb1a* | -0.68 | 0.003 | **0.024** |
|  | *Abcg2* | 0.51 | 0.09 | 0.21 |
| *Bacillus* | *Abcb1a* | -0.61 | 5.3 x 10<sup>-6</sup> | **0.0005** |
|  | *Abcg2* | 0.75 | 0.002 | **0.023** |
| *Eubacterium* | *Abcb1a* | -0.67 | 0.002 | **0.023** |
|  | *Abcg2* | 0.79 | 0.002 | **0.023** |
| *Xiphinematobacter* | *Abcb1a* | -0.79 | 0.002 | **0.023** |
|  | *Abcg2* | 0.78 | 0.01 | 0.059 |
| *Clostridium* | *Abcb1a* | -0.78 | 0.0003 | **0.009** |
|  | *Abcg2* | 0.65 | 0.05 | 0.15 |
| *Legionella* | *Abcb1a* | -0.83 | 0.002 | **0.023** |
|  | *Abcg2* | 0.69 | 0.11 | 0.23 |
| *Methanogenium* | *Abcb1a* | 0.73 | 0.005 | **0.04** |
|  | *Abcg2* | -0.63 | 0.07 | 0.19 |
| *Acholeplasma* | *Abcb1a* | 0.66 | 0.002 | **0.023** |
|  | *Abcg2* | -0.92 | 5.1 x 10<sup>-5</sup> | **0.003** |
| *Pseudanabaena* | *Abcb1a* | 0.80 | 0.001 | **0.023** |
|  | *Abcg2* | -0.85 | 0.002 | **0.023** |
| *Anaeroplasma* | *Abcb1a* | 0.74 | 0.003 | **0.024** |
|  | *Abcg2* | -0.75 | 0.008 | 0.057 |

Refer to heatmap Figure 7B.

## Supplementary Figure A.1

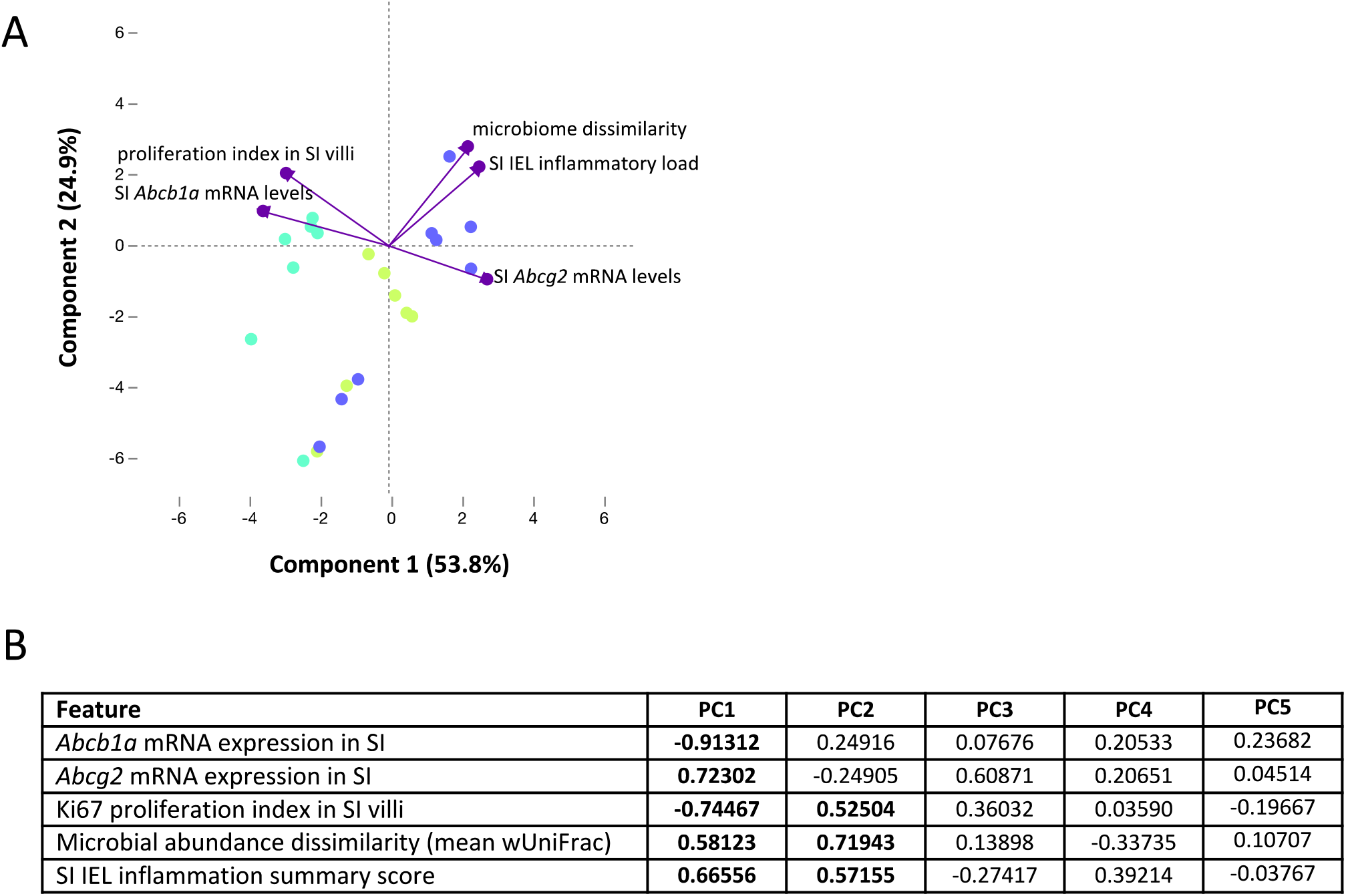

